# Dendritic speeding of synaptic potentials in an auditory brainstem principal neuron

**DOI:** 10.1101/688309

**Authors:** Geetha Srinivasan, Andre Dagostin, Richardson N. Leão, Veeramuthu Balakrishnan, Paul Holcomb, Dakota Jackson, George Spirou, Henrique von Gersdorff

**Author notes:** **Corresponding author:** Henrique von Gersdorff, PhD.

## Abstract

Principal cells of the medial nucleus of the trapezoid body (MNTB) in the mammalian auditory brainstem receive most of their strong synaptic inputs directly on the cell soma. However, these neurons also grow extensive dendrites during the first four postnatal weeks. What are the functional roles of these dendrites? We studied the morphology and growth of the dendrites in the mouse MNTB using both electron microscopy and confocal fluorescence imaging from postnatal day 9 (P9; pre-hearing) to P30. The soma of principal cells sprouted 1 to 3 thin dendrites (diameter ~ 1.5 microns) by P21 to P30. Each dendrite bifurcated into 2-3 branches and spanned an overall distance of about 80 to 200 microns. By contrast, at P9-11 the soma had 1 to 2 dendrites that extended for only 25 microns on average. Patch clamp experiments revealed that the growth of dendrites during development correlates with a progressive decrease in the input resistance, whereas acute removal of dendrites during brain slicing leads to higher input resistances. Accordingly, recordings of excitatory postsynaptic potentials (EPSPs) evoked by afferent fiber stimulation show that EPSP decay is faster in P21-24 neurons with intact dendrites than in neurons without dendrites. This dendritic speeding of the EPSP reduces the decay time constant 5-fold, which will impact significantly synaptic current summation and the ability to fire high-frequency spike trains. These data suggest a novel role for dendrites in auditory brainstem neurons: the speeding of EPSPs for faster and more precise output signal transfer.

**Significance Statement:** Auditory circuits that compute sound localization express different types of specialized synapses. Some are capable of fast, precise and sustained synaptic transmission. As the paradigm example, principal cells of the MNTB receive a single calyx-type nerve terminal on their soma and this large excitatory synapse produces fast and brief supra-threshold EPSPs that can trigger trains of high frequency spikes. However, these neurons also extend thin and long dendrites with unknown function. We examined the relationship between dendritic morphology, passive electrical properties and EPSP waveform. We found that more mature neurons with intact dendrites have lower input resistances and short EPSP waveforms, ideally suited for conveying precise timing information, whereas immature neurons with shorter dendrites and higher input resistance have longer lasting EPSPs.

## Introduction

Neurons communicate with each other via synapses that are mostly located on extended and highly branched dendritic trees (Reyes, 2001; Martina et al., 2003). The complexity of synaptic integration arises from a wide variation in neuronal soma, axonal and dendritic morphology, ion channel expression and location, and how these factors are developmentally regulated during action potential firing (Nicoll et al., 1993; Libersat and Duch, 2004; Gulledge et al., 2005). Indeed, the shape of EPSPs can be influenced by the presence of an axonal current sink, an effect termed axonal speeding of the EPSP (Mejia-Gervacio et al., 2007). Moreover, amputation of dendrites in cerebellar Purkinje and cortical pyramidal neurons leads to an increase in input resistance and changes in neuronal excitability (Bekkers and Hausser, 2007). The morphology and membrane properties of axons, somas and dendrites can thus sculpt the EPSP waveform.

The time course of the EPSP is determined by the waveform of the depolarizing EPSC and the intrinsic membrane properties. The currents active near the resting potential of brainstem auditory neurons include voltage-independent potassium leak currents (e.g. two-pore K channels; Bernston and Walmsley, 2008), voltage-gated low-threshold potassium current (I_KLT_; Manis and Marx, 1991; Brew and Forsythe, 1995) and hyperpolarization-activated cation currents (I_h_; Golding et al., 1995). Together these currents determine the input resistance (R_i_) and contribute to the membrane time constant (*τ*_M_), which shapes the neuron’s voltage response to the EPSC. Specialized time-coding auditory brainstem neurons can have very low R_i_ and fast *τ*_M_ (Scott et al., 2005). In fact, some avian and mammalian neurons can have an extremely fast EPSP time course that is nearly identical to that of the EPSC (MacLeod and Carr, 2012).

The medial nucleus of the trapezoid body (MNTB) is one of the several nuclei in the superior olivary complex of mammals. MNTB neurons fire at very high frequencies and phase-lock to the presynaptic input or to its temporal envelope (Spirou et al., 1990; Wu and Kelly, 1993; von Gersdorff & Borst 2002; Tollin, 2003). The MNTB is classically thought to function as a relay station between the cochlear nucleus (CN) and the lateral and medial superior olives, playing a major role among those brainstem nuclei that are involved in binaural hearing (Kandler and Friauf, 1993; Smith et al., 1998). The thick caliber and myelinated axons of the globular bushy cells of the anterior ventral CN form giant synaptic terminals called calyces of Held on the soma of MNTB principal cells (Morest, 1973; Kuwabara et al., 1991). The calyx of Held covers a large fraction of the MNTB cell soma where it forms multiple heterogeneous active zones (Rowland et al., 2000; Sätzler et al., 2002; Hoffpauir et al., 2006; Spirou et al., 2008). The large number of active zones results in large amplitude EPSCs that ensure rapid and secure synaptic transmission during repetitive afferent fiber stimulation (Taschenberger et al., 2002; Lorteije et al., 2009; Fekete et al., 2019), although synaptic inhibition can cause spike failures (Kopp-Scheinpflug et al., 2011).

The adult MNTB principal cell also has extensive dendrites (Banks and Smith, 1992; Sommer et al. 1993), which contain Na^+^ and K^+^ channels that contribute to neuronal excitability (Leão et al., 2008; Elezgarai et al., 2003). How dendritic properties change during early postnatal development and whether these properties contribute to synaptic integration remains poorly understood in the MNTB. Here we study the morphology of P9 to P24 MNTB dendrites using confocal microscopy and we compare this to more mature P30 neurons using EM reconstructions. We show that immature P9-11 MNTB neurons have shorter dendrites, smaller resting membrane capacitance and higher input resistance than P21-28 neurons that extend long and thin dendrites. Importantly, we found that more mature P21-24 MNTB neurons with intact dendrites have significantly faster EPSP decay kinetics than P21-24 neurons without long dendrites. Our computer modeling shows that a long and thin dendrite with a large leak conductance can produce faster EPSP decays. We therefore propose that MNTB neurons sprout a dendritic tree during early postnatal development in part to reduce input resistance and decrease their membrane time constant and, thus, speed EPSP decay times.

## Materials and Methods

All experiments were performed according to protocols approved by the Oregon Health and Science University and University of West Virginia IACUC (Institutional Animal Care and Use Committee) in accordance with NIH guidelines.

### Brainstem slice preparation

Acute brainstem slices were prepared from postnatal day 5 (P5) to P24 mice pups of either sex (C57BL/6J strain; Charles River Laboratories, Wilmington, MA). After decapitation, the brainstem was quickly immersed in ice-cold low calcium artificial cerebrospinal fluid (aCSF) containing (in mM): 125 NaCl, 2.5 KCl, 3 MgCl_2_, 0.1 CaCl_2_, 25 glucose, 25 NaHCO_3_, 1.25 NaH_2_PO_4_, 0.4 ascorbic acid, 3 myo-inositol, 2 Na-pyruvate, pH 7.4-7.5 when bubbled with carbogen (95% O_2_, 5% CO_2_) and having osmolarity of 310-320 mOsm. For older animals (P21-24), the entire procedure was performed in room temperature. Transverse slices of the auditory brainstem were cut at a thickness of 200 μm using a vibratome slicer (VT1000; Leica, Bannockburn, IL), and incubated at 37°C for 30 min in normal aCSF and thereafter kept at room temperature (22-24°C) for experiments. The normal aCSF was the same as the low-calcium aCSF except that 1 mM MgCl_2_ and 1.2 or 2 mM CaCl_2_ were used.

### Patch-clamp electrophysiology

After incubation, slices were transferred to a 1 ml chamber and perfused with normal aCSF at the rate of 1.5-2 ml/min. Medial nucleus of the trapezoid body (MNTB) neurons were viewed using a Zeiss Axioskop 2 FS microscope equipped with differential interference contrast (DIC), and a 40x water-immersion objective. To perform whole-cell recordings from MNTB neurons, we used an intracellular solution containing the following (in mM): 130 K-gluconate, 20 KCl, 5 Na_2_-phosphocreatine, 10 HEPES, 5 EGTA, 4 Mg-ATP, and 0.5 GTP, pH adjusted to 7.3 with KOH, and 300-305 mOsm. In addition, 4 mM QX-314 was added in the internal solution to measure excitatory postsynaptic currents (EPSCs) and potentials (EPSPs). All salts were purchased from Sigma (St. Louis, MO).

Recording pipettes were pulled from borosilicate glass capillaries (World Precision Instruments) with a Sutter P-97 electrode puller (Sutter Instruments, Novato, CA) and had open tip resistances of 2.0-3.0 MΩ. Access resistance (R_s_ was ≤ 9 MΩ and R_s_ was compensated >85%, so that the compensated R_s_ was about 1-2 MΩ. All traces for kinetic analysis and display were corrected off-line for series resistance and holding potential errors (Schneggenburger et al. 1999). For voltage-clamp recordings, the principal cells were voltage clamped at holding potential of −70 mV and the synaptic signals were filtered at 2.9 kHz. For analysis of passive membrane properties, capacitive currents elicited by 10 mV hyperpolarizing voltage step were filtered at the minimum amount (filter 1 at 30 kHz and filter 2 at 13.9 kHz) and the sampling rate was 100 kHz. Afferent fibers were stimulated with a bipolar platinum/iridium electrode (Frederick Haer Company, Bowdoinham, ME) placed near the midline spanning the afferent fiber tract of the MNTB. An Iso-Flex stimulator driven by a Master 8 pulse generator (A.M.P.I., Jerusalem, Israel) was used to deliver step pulses (100 μs,<15 V DC). Except capacitive currents (see above) the voltage-clamp and current-clamp data were acquired at 10 or 25 μs sampling rate using an EPC-9/2 or EPC-10/2 amplifier (HEKA Elektronik, Lambrecht, Germany) controlled by Pulse 8.4 or PatchMaster software and filtered on-line at 2.9 kHz.

### Confocal fluorescence microscopy

MNTB neurons were filled with Alexa fluor 555 (250 μM) via the patch pipette for at least 20 min. The 200 μm thick brainstem slices were subsequently fixed in 4% (wt/vol) paraformaldehyde in phosphate buffer solution (PBS) overnight at 4°C. Slices were then washed in PBS three times, and each wash lasted about 15 min, and the slices were then mounted onto superfrost slides in photobleaching-protective medium (Gel/Mount™, biomeda corp). Stained slices were viewed with laser lines at 543 nm (orange red) using a 20x objective on a confocal-scanning microscope (Olympus Fluoview 300). The confocal images were analyzed using ImageJ (Wayne Rasband, NIH).

### Data Analysis

Data were analyzed off-line and presented using Igor Pro (Wavemetrics, Lake Oswego, OR) and AxographX. Results are expressed as mean ± S.E.M. The significance of differences among data sets was evaluated by Student’s unpaired two-tailed *t* test; p<0.05 was considered as significant.

### Estimation of decay kinetics, capacitance and input resistance via voltage-clamp

Capacitive currents elicited by a voltage-clamp step of 10 mV were fitted over a time interval of 25 ms with mono, bi, or tri exponential function (based on the improvement of the summed square error):

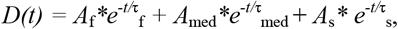

where *D(t)* is the decay of the current as a function of time (*t*) and *A*_f_, *A*_med_ and *A*_s_ are amplitude constants and *τ*_f_, *τ*_med_, and *τ*_s_ are the fast, medium, and slow decay time constants, respectively. We first subtracted any capacitive currents in the cell-attached mode to eliminate the current due to the pipette capacitance and seal resistance. The time point used to estimate the initial amplitude of capacitive currents was taken at 0.1 ms after the start of the voltage-clamp jump (see Methods section of Mejia-Gervacio et al., 2007 and Nadeau and Lester, 2000; and appendix of Nadeau and Lester, 2002). The amplitudes and the time constants of the capacitive current are A_f_, A_med_, As, and τ_f_, τ_med_, τ_s_ respectively. Individual capacitance components were estimated from:

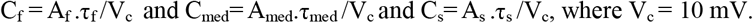

The cell input resistance R_I_ = V_c_/(A_f_ + A_med_ + A_s_) and the steady-state input resistance was R_Iss_ = V_c_/I_ss_, where I_ss_= steady-state capacitive current. Membrane capacitance (C_m_) was also calculated from C_m_= C_sp_ x S, where C_sp_ = specific capacitance (10 fF/μm^2^) and S = surface area. The decays of the evoked excitatory postsynaptic currents and potentials (EPSCs and EPSPs) were fit by single- or double-exponential functions. The weighted decay time constant of the EPSC or EPSP was calculated as

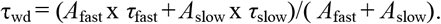

### Computer modeling

Based on our anatomical data, we implemented a multi-compartment model of the MNTB principal neuron to investigate the contribution of different structural compartments to synaptic integration using the *NEURON* simulation environment (Hines & Carnevale, 2001; for more details of the specific model see Leão et al., 2008). MNTB neurons were modeled with no dendrites, or one dendrite compartment with various lengths and the diameters (Ø) set to 3 μm emerging from a spherical soma (Ø = 20 μm). We also included a cylindrical axonal compartment originating at the soma (L = 20 μm, Ø = 2 μm). Each dendritic (and axonal) cylinder was divided into 10 isopotential compartments connected by axial resistances (150 Ω/cm). All the compartments were passive and we implemented various leak conductances with reversal potentials equal to the resting potential of the cell (−65 mV). In addition, we implemented a single short EPSP provided by a calyceal input connected to the MNTB soma using the calyx of Held model of Graham et al (2001) and Hennig et al., 2008.

### EM segmentation and 3D Reconstruction from SBEM

Segmentation of MNTB principal cells, dendrites and axons was performed on a serial block-face scanning electron microscopy (SBEM) image volume collected by the National Center for Microscopy and Imaging Research (NCMIR, University of California: San Diego) at P30. This image volume measured 154.81μm x 77.51μm x 142.3μm with pixel dimension of 4 nm in X and Y, and a slice thickness of 50 nm. The image volume was taken from the medial portion of the MNTB. Segmentation was accomplished by hand using the paint brush tool in the Seg3D software (Scientific Computing and Imaging Institute, University of Utah) and 3D reconstruction was performed using the ISO command to produce isosurfaces, which were exported as VTK objects. For maximum compatibility with other software, these VTK objects were converted to OBJ format using a custom Python script. All images of 3D reconstructions were rendered using the Rhinoceros 3D software (Robert McNeel and Associates, Seattle, WA).

### Dendrite length and diameter measurement

Dendrites from SBEM images were skeletonized manually using their 3D reconstructions in the syGlass virtual reality software (IstoVisio, Morgantown, WV) by placing points in the center of the reconstruction at regular intervals. Skeletons were exported in SWC format and converted to comma-separated value files (CSV) using a custom Python script. The CSV files were imported along with their respective 3D reconstructions into Rhinoceros 3D for further analysis. Polylines representing the dendrite skeleton were created using the CurveThroughPoint command, with some manual editing to separate branching paths. The resulting branch lengths were measured using the Length command, and then broken into 10 μm segments beginning at the most proximal point using the Divide command. Four diameter measurements were taken at each 10μm interval by creating 4 lines perpendicular to the dendrite skeleton and at 45° to one another and trimming them by the dendrite OBJ. These measurements were then averaged, resulting in one measurement at each 10 μm interval. All graphs were generated using Matlab (MathWorks, Natick, Massachusetts).

## Results

### Mouse MNTB principal cell serial block-face electron microscopy

At the end of the fourth postnatal week the mouse auditory system is adult-like by several functional measures (Liberman and Liberman, 2016). Using serial block-face scanning electron microscopy (SBEM) we thus first determined the *in situ* dendritic morphology of P30 mouse MNTB neurons as a benchmark of adult-like neuronal morphology. We reconstructed a total of 8 principal neurons in the medial MNTB (see Materials and Methods for SBEM procedures; **Figure 1A**). Each neuron had one to three dendrites sprouting from the cell soma with a few (2-4) secondary branches and a fairly constant diameter of about 1.5 microns. Most of the myelinated axons aligned with one another as they exited the MNTB forming a fiber bundle (right hand side of Figure 1A). Two isolated cells (**Figure 1B and 1C**; white arrows) illustrate the length, thinness, and sparsity of the dendrites and their lack of profuse dendritic branching or spines. Although the main excitatory drive of the MNTB principal cell, the calyx of Held synapse, is located entirely on the cell soma, these data clearly show that dendrites remain a stable and prominent structure that is not pruned away during early development. Given the large calyx terminal on the principal cell soma, which drives large EPSPs and spikes, the functional role of dendrites becomes an important question.

**Figure 1.**
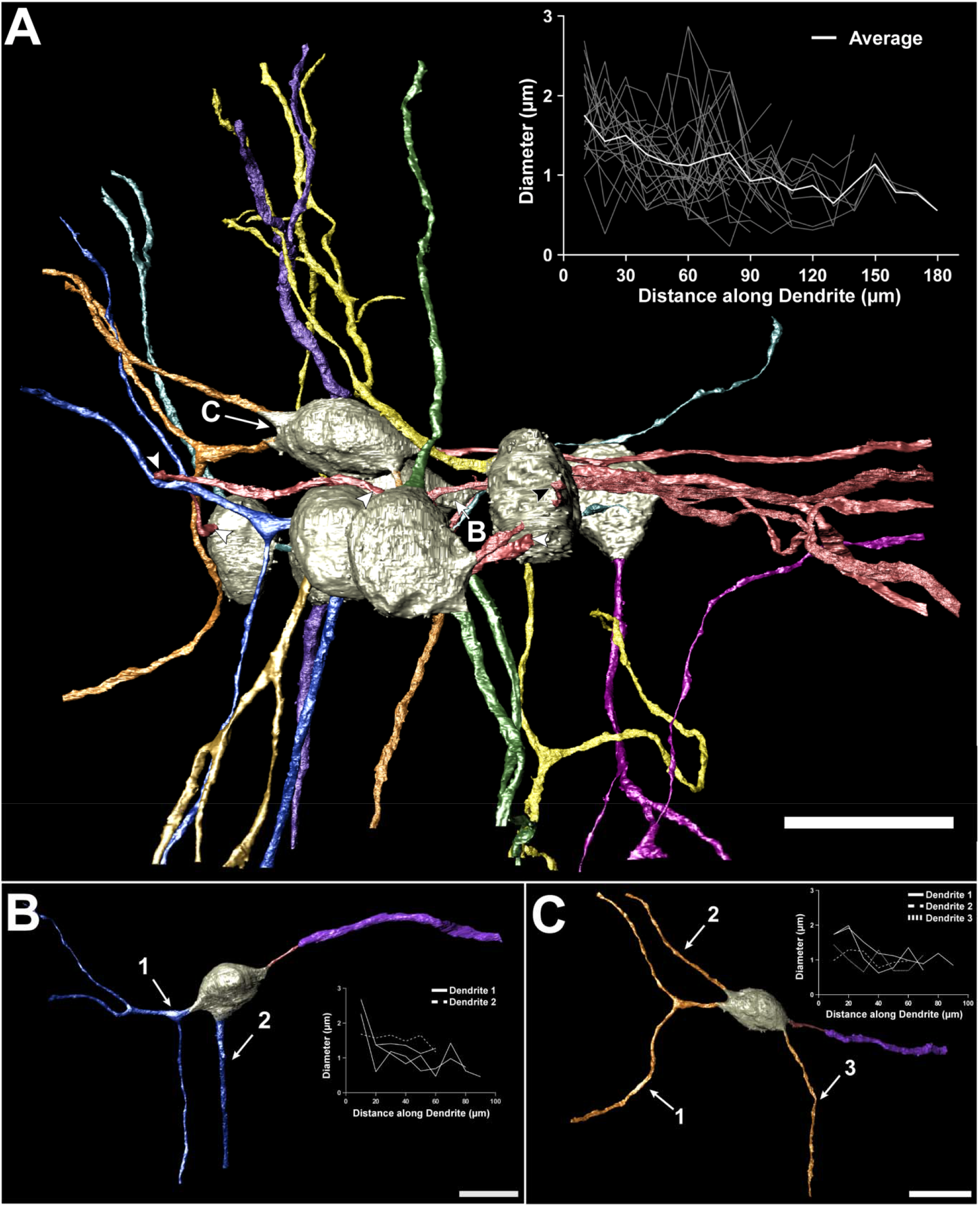
3D reconstruction of mouse MNTB principal cells. (**A**) Eight MNTB cells from postnatal day P30 reconstructed from a serial block-face scanning electron microscopy image volume, including cell bodies (tan), dendrites (multiple colors), and axons (rose). The dendrites (1-3 per cell) tend to be long and straight, with relatively few branches and a fairly constant diameter (A, inset graph). Dendrites proximal to one another exhibit similar orientations, as do the MTNB cell axons. The axon from the left-most cell takes a tortuous path (white arrowheads) before it aligns with the axon of a neighboring cell and is clipped at the edge of the image volume. A third axon (black arrowhead) appears to take a similar trajectory to the first two axons, but is also quickly clipped at the edge of the image volume as well. The remaining five axons align with one another and take a similar path exiting the image volume to the right, forming an axon bundle. (**B, C**) Two cells (white arrows) isolated in the lower panels highlight the sparsity of dendrites and lack of dendritic branches or spines. Additionally, myelin (purple) can be seen ensheathing the axons of each of the cells, revealing that the axon initial segment is bare of myelin. Scale bars = 20 μm.

We used our reconstruction of the MNTB cells to calculate the total cellular surface area and the individual contributions to this area of the cell soma, dendrites and axons within our reconstructed volume. The total average area was about 3482 μm^2^ if we include the full axon area and 3043 μm^2^ if we include just the axon initial segment (**Table 1**). About 44% of this was the soma, 40% was the dendrites, and ~16% was the thin axons. The dendrites thus contributed a significant portion of the cell area. Since some of the dendrites were cut off, this is an underestimate of the dendritic contribution. These results allowed us to estimate the cell membrane capacitance (C_m_) using the specific capacitance of lipid bilayer membranes (9 to 10 fF/μm^2^; Gentet et al., 2000). The total C_m_ of the MNTB principal cell is thus between 31.3 pF to 34.8 pF (Table 1). Due to fixation and dehydration for EM processing we estimate that the cell surface area may shrink by 20% (see Kuba et al., 2005). Thus, the *in situ* living cell C_m_ may be about 38 to 41 pF. However, myelin greatly reduces the axonal capacitance. The mature MNTB cell capacitance (with just axon initial segment) may thus be closer to 27.4 to 30.4 pF or, after the 20% correction for shrinkage, 33 to 36 pF.

**Table 1.**
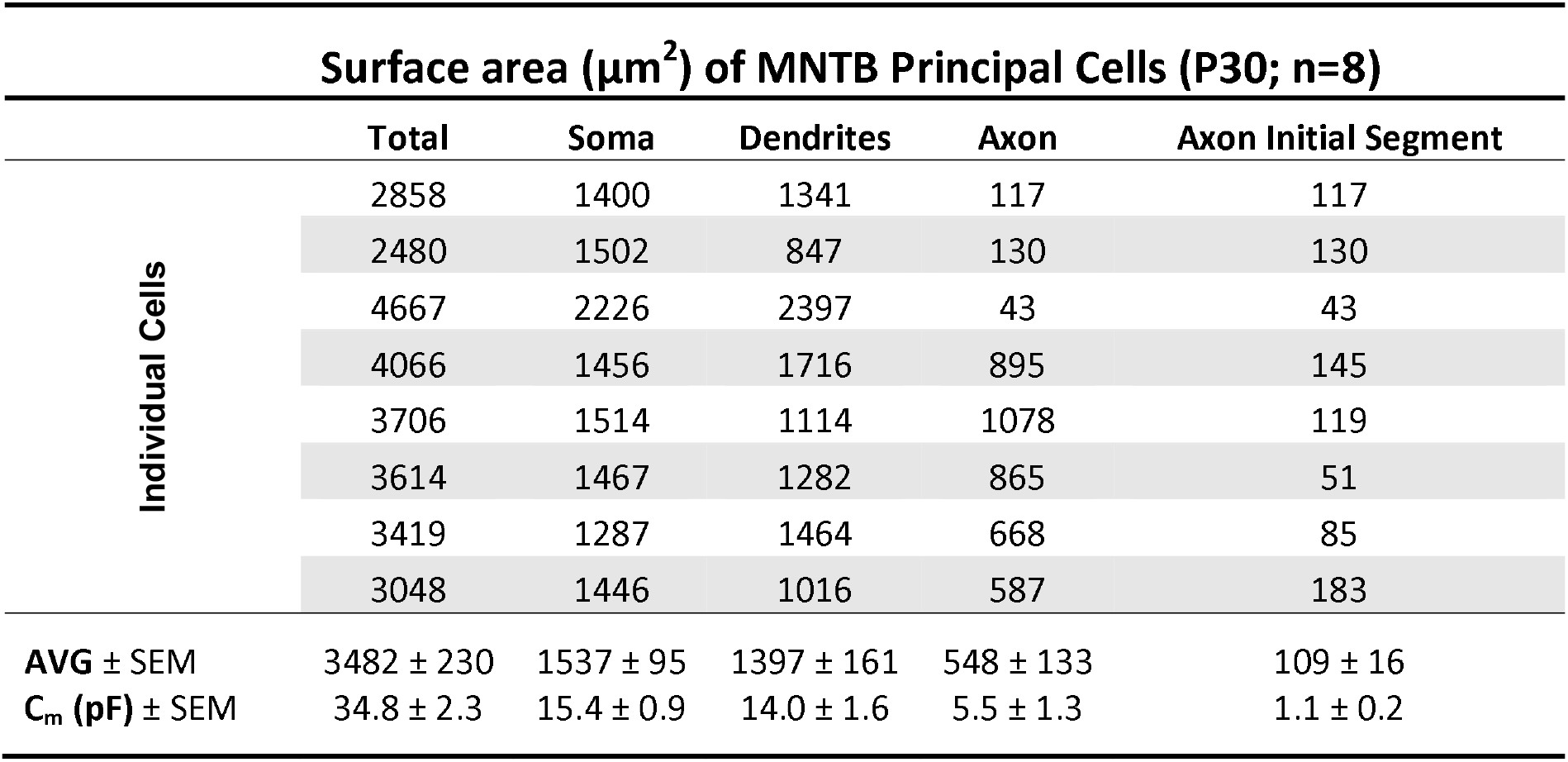
MNTB principal cell surface area and membrane capacitance at postnatal day 30 (P30). Data from 8 different cells is shown for the total surface area and for the separate surface areas of soma, dendrites and axon, as well as the axon initial segment.

We also determined the number and location of synaptic boutons on the dendrites (**Figure 2A**). Synaptic inputs were sparse along the dendrites and clustered near the initial emergence of the dendrite. The number of dendritic inputs falls off as a function of distance from the soma (n=16 dendrites from 8 MNTB cells; **Figure 2B**). The dendritic membrane not innervated was often contacted and covered by thin glial processes (**Figure 2C and 2E**). The dendrite was sometimes completely ensheathed by glial processes, which could be buffering external potassium efflux from the dendrite (Elezgarai et al., 2003).

**Figure 2:**
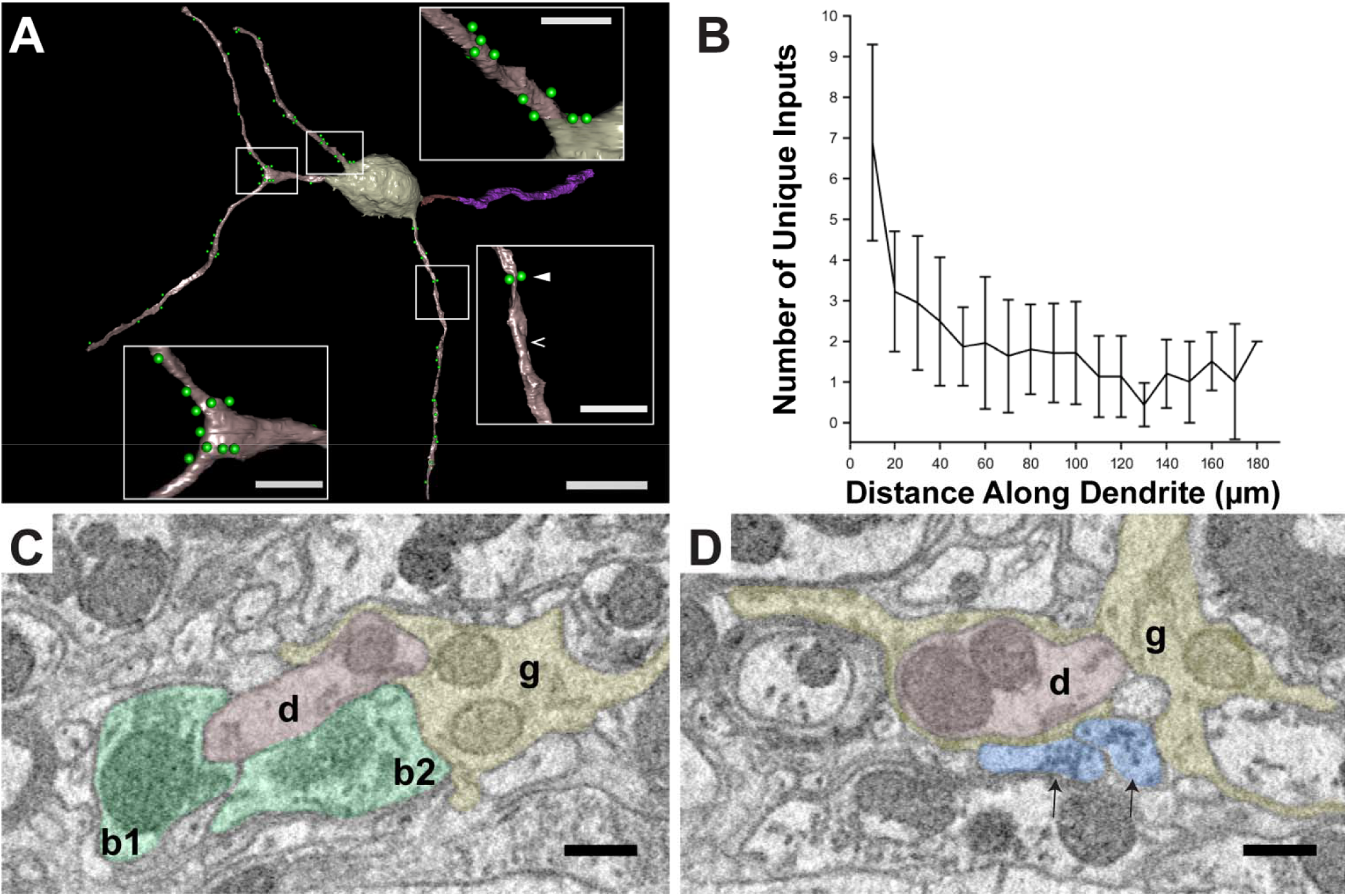
Dendritic inputs are dense proximal to the MNTB soma and become sparse at more distal locations. (**A**) MNTB principal cell at P30 with soma (tan), axon with myelin (maroon and purple, respectively), and 3 thin dendrites (brown), with only one showing a bifurcation and no distinct dendritic spines. Inputs (green spheres) are found clustered either at dendritic swellings (lower left inset) or at the initial emergence of the dendrite from the soma (upper inset). Inputs along the thin main dendritic body are sparse (lower right inset). Panels C and D represent electron microscopy images taken from the locations in the lower right panel denoted by the white filled arrowhead and open arrowhead, respectively. (**B**) The number of unique dendritic inputs falls off quickly as a function of distance from the soma along the dendrite (n=16 dendrites from 8 unique MNTB cells). (**C**) EM of location along dendrite in (A) denoted in lower right panel by filled white arrowhead. The dendrite (d, brown) is innervated by two distinct boutons (b1 and b2, green) filled with round vesicles. The dendritic membrane not innervated is covered by thin glial processes extending from a glial projection (g, yellow). (**D**) EM of location along dendrite in (A) denoted in lower right panel by open white arrowhead. The dendrite (d) in this location is completely ensheathed by glial processes (g, yellow), preventing two potential boutons (blue) from making contact. Scale bars: A, 20μm; A (insets), 5μm. C-D, 0.5μm.

### Input resistance and membrane time constant of MNTB neurons

Previous morphological reconstructions of MNTB principal cells with neurobiotin and HRP-injection reveal one to three dendrites that sometimes extended beyond the borders of the MNTB (3-5 week-old rats, Banks and Smith, 1992; adult rats, Sommer et al., 1993; P16 rats, Leão et al., 2008). One or two primary dendrites sprung from the soma and then bifurcated into several smaller-diameter, aspiny branches. The axon projected mostly to SPN, MSO and LSO. The input membrane resistance (R_m_) of the cells measured with sharp conventional electrodes and hyperpolarizing currents was about 65 MΩ. Subsequent, patch clamp recordings in gerbil MNTB revealed an R_m_ that declined with age from 130 MΩ at P14-16 to 115 MΩ at P19-20 to 86 MΩ at P28-29 (Scott et al., 2005). The membrane time constant (*τ*_M_) was found to be stable during this developmental period (*τ*_M_ = 3.6 to 4.3 ms). Biotin-filled MNTB neurons revealed one to two small caliber primary dendrites (Scott et al., 2005).

Here we calculated R_m_ using first current-clamp recordings from mouse MNTB neurons. **Figure 3A** shows recordings from a P20 mouse MNTB principal cell at room temperature. Cells were held at a resting membrane potential of −70 mV. Depolarizing step current injections of 150 pA elicited one action potential spike and outward rectification, consistent with previous reports (Banks and Smith, 1992; Forsythe and Barnes-Davies, 1993; Wang et al., 1998). The spike threshold was about −45 mV. The IV curve was generated using 200 ms current pulses from −300 pA to +150 pA relative to the holding current. The pulses were applied in 50 pA steps with 3 s inter pulse interval. The IV curve were built with values obtained from the peak of the hyperpolarizing sag (black dots) and the mean of the last 10 ms (open triangle; **Figure 3A**). The average IV curve was linear (ohmic; dashed line) over a range of negative current with a slope (input resistance) R_m_ = 207 MΩ (**Figure 3B**). Deviations from linearity are due to activation of I_KLT_ for positive currents and I_h_ for negative currents (Brew and Forsythe, 1995; MacLeod and Carr, 2012). For small current injections, the average R_m_ = 247 ± 65 MΩ (n=6; P20). This was not significantly different from P28 neurons: R_m_= 200 ±21 MΩ (n=4; P28; 24°C). This was however substantially larger than R_m_ for P20 gerbils (Scott et al., 2005) or 3-5 week old rats (Banks and Smith, 1992). We attribute this difference, in part, to the smaller size of mouse MNTB neurons, compared to gerbil or rat neurons, and to room temperature (see Figure 3E).

**Figure 3.**
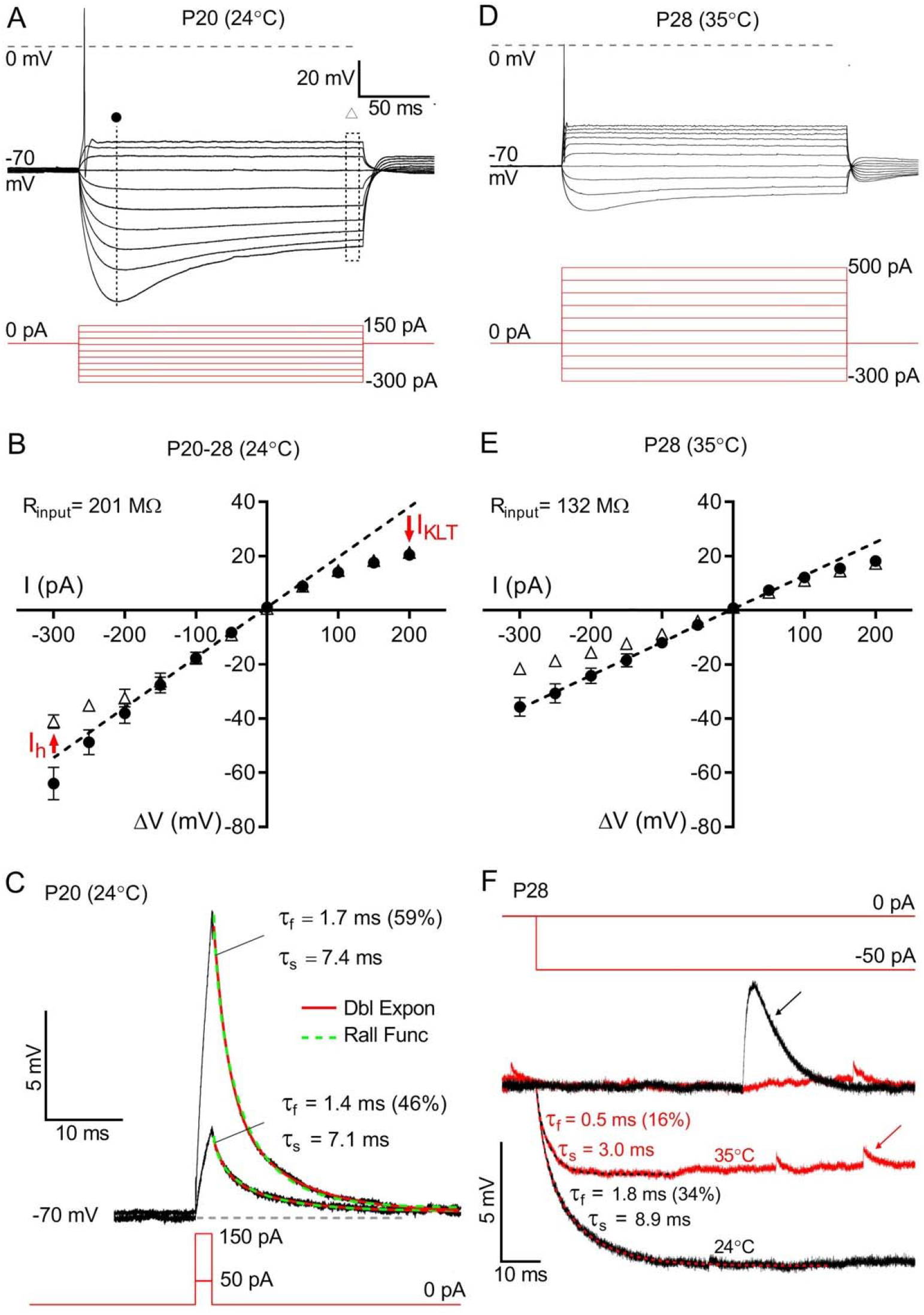
Current clamp responses of MNTB principal cells. (**A,B**) The voltage responses (black) recorded from a P20 neuron after step current injections (red; 50 pA steps) at 24°C. At the threshold a single action potential spike is triggered by a step depolarization. The vertical dotted line marks the membrane potential values at the onset of the I_h_ current plotted as closed circles in panel B and the dotted rectangle shows the time period over which the steady-state membrane potential was measured (open triangles in panel B; values plotted as mean ± SEM, n=6). The slope of the dashed line in panel B (the I-V curve) is a measure of the average input resistance of the cell (R_m_ = 201 MΩ). The deviation from a straight line at depolarized potentials is due to the activation of a low threshold potassium current (I_KLT_) and at hyperpolarized potentials is due to the activation of I_h_. (**C**) A 2 ms current injection (+50 and +150 pA in red, bottom) generates rapid EPSP-like voltage responses (black). The decay of the transient voltage response is well fit by a double exponential function (superimposed red trace). The respective fast and slow time constants, and the percent contribution of the fast component, are shown. The same data can also be well fit by a Rall function (dashed green curve). (**D**) The voltage responses from a P28 neuron after step current injections (red; 100 pA steps) at 35°C. At the threshold a single action potential spike is triggered by a step depolarization. (**E**) Same as in panel B but for P28 neurons at 35°C. The slope of the dashed line (R_m_ =132 MΩ) is lower than for panel B. This decrease of input resistance is also evident in panel D. (**F**) The effect of temperature on membrane passive properties. Current clamp recordings from the same cell in response to a step current injection (−50 pA; black trace, 24°C; red, 35°C). The dashed superimposed lines are best fits to a double exponential function with the respective fast (τ_f_) and slow (τ_s_) time constants shown on the right. The percent contribution of the fast component is also shown. The mean membrane time constant is faster at 35°C. Arrows indicate spontaneous postsynaptic currents (PSC) in both traces.

We found that a short 2-ms current injection elicits a voltage transient that decays as a double-exponential function (**Figure 3C**). The fast (τ_f_) and slow (τ_s_) components of the membrane time constant (τ_m_) were unchanged for different depolarizing steps (50 pA: 1.6±0.5 and 6.5±0.7 ms; 150 pA: 1.3±0.4 and 6.3±0.7ms for τ_f_ and τ_s_, respectively; n=4; **Figure 3C**). This double-exponential decay of the EPSP suggests that the MNTB cell cannot be modeled as a passive isopotential spherical cell or point-neuron, which would have only a single exponential decay with τ_M_ = C_m_R_m_ (Golding, 2012). If we assume a “stick-and-ball” passive neuron model the voltage V(t) as a function of time should decay according to the Rall function (Rall, 1969):

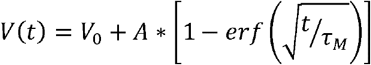

where V_0_ and A are constants and *erf*(x) is the error function:

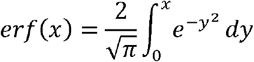

The best fit values for τ_M_ are 6.4 ± 0.7 ms (n=6) and 7.8 ± 0.8 ms (n=5) for 50 and 150 pA current pulses respectively, which are not significantly different (unpaired Student t test; p>0.05). As shown in Figure 3C both a double exponential function and a Rall function provide excellent fits to the data.

**Figure 3D** shows current clamp responses of more mature neurons (P28) at more physiological temperatures (35°C). To elicit an AP spike larger step current injections are now required than for room temperature. This is due to a significant reduction in the input resistance of the neurons at high temperature (for the same P28 neurons: R_m_ = 200 ± 21 MΩ at 24°C and R_m_ = 119 ± 19 MΩ at 35°C; n=4; p=0.0002 paired Students t-test; **Figure 3E**). Current injection in the soma of a nonisopotential neuron leads to current flow between different compartments and V(t) can be expressed as the sum of exponential functions (Rall, 1969; Major et al., 1993). **Figure 3F** shows voltage responses of a P28 neuron to small hyperpolarizing pulses (−50 pA step) for two temperatures. The responses again require a double exponential fit with both fast (τ_f_) and slow (τ_s_) time constants. The value of τ_s_ = τ_M_ (the membrane time constant) and τ_f_ is related to the dendrites and axon (Rall, 1969). The electrotonic length is 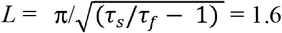 for P20 and P28 MNTB neurons at 24°C (n=4; Rall, 1969), which is larger than values for hippocampal neurons (*L* = 0.9; Brown et al., 1981), which have thicker dendrites (9-10 μm). The total membrane capacitance can be calculated using C_m_= τ_s_ /R_0_, where R_0_ is the input resistance times the percentage of the amplitude of the slow component (59 ± 5%; n=6; see Golowasch et al., 2009; Major et al., 1993). The average C_m_ calculated in this manner is 45.0 ± 4.5 pF (n=6 for 24°C).

### Developmental changes in passive membrane properties of MNTB neurons

Capacitive currents from passive and multi-compartment neurons can be used to estimate the C_m_ of the different compartments under voltage clamp (Mejia-Gervacio et al., 2007; Nadeau and Lester, 2000 and 2002). This analysis reveals three main compartments: soma, axon and dendrites. To better correlate changes in neuronal compartments with our estimates of C_m_ we performed measurements at four different age groups (P5-6, P9-11, P15-17 and P21-24). Capacitive currents were measured in response to −10 mV hyperpolarizing voltage steps, immediately after break in, with minimum filtering. The decay time course of the capacitive current was fit with exponential functions. In the youngest group (P5-6), capacitive current transients were well-fitted with the sum of two exponential components which we shall refer to as the fast and medium components, respectively (**Table 2**). Likewise, in P9-11 animals, 87% of MNTB neurons exhibited two exponential decay components and 10% exhibited a single component. However, the remaining 3% exhibited a third, much slower component. In the oldest group (P21-24) the percentage of neurons with a tri-exponential decay increased significantly to 46% (**Figure 4**). In addition, the steady-state input resistance (R_I_) significantly dropped during post-natal development, from 0.8±0.1 GΩ at P5-6 (n=5) to 0.6±0.06 GΩ at P9-11 (n=26) and then to 0.2±0.03 GΩ for P21-24 neurons (n=16; p=0.0002) with triexponential decay. Steady-state input resistance in P21-24 neurons with bi-exponential decay was higher: 0.5±0.09 GΩ (n=19). Membrane input resistances at 24°C were similar when we performed voltage and current clamp recordings in the same group of cells (P20: 246 ± 60 and 250 ± 70 MΩ, respectively; n=6). We conclude that during the first three weeks the passive properties of mouse MNTB neurons undergo substantial changes, but are not significantly different from P20 to P28.

**Table 2.** Capacitive current decay time constants and input resistance during postnatal development. The % of cells^**a**^ at different postnatal ages: At P5-6, 100% cells showed bi-exponential decay. At P9-11 (n=30), 87% cells showed bi-exponential decay, 10% showed mono-exponential and 3% showed triexponential. At P15-17 (n=14), 71% cells showed bi-exponential decay, 7% showed mono-exponential and 22% showed triexponential. At P21-24 (n=35), 54% cells showed bi-exponential decay, and 46% showed tri-exponential.

| Postnatal age | $\tau_f$ (ms) | $\tau_m$ (ms) | $\tau_s$ (ms) | C<sub>f</sub> (pF) | C<sub>m</sub> (pF) | C<sub>s</sub> (pF) | A<sub>f</sub> (pA) | A<sub>m</sub> (pA) | A<sub>s</sub> (pA) | % of cells<sup>a</sup> | R<sub>i</sub> (GΩ) |
| --- | --- | --- | --- | --- | --- | --- | --- | --- | --- | --- | --- |
| **P5-6** (n=5) | 0.15±0.04 | 2.0±0.30 | | 10±1.4 | 14±1.5 | | 800±120 | 63±11 | | 100% | 0.8±0.1 |
| **P9-11** (n=26) | 0.2±0.01 | 2.0±0.15 | | 11±0.7 | 14±1.5 | | 671±49 | 78±8 | | 87% | 0.6±0.06 |
| **P15-17** (n=10) | 0.2±0.02 | 2.1±0.4 | | 9.64±1.0 | 15±2.0 | | 669±80 | 78±15 | | 71% | 0.6±0.1 |
| **P21-24** (n=16) | 0.15±0.01 | 1.6±0.2 | 9.6±1.3 | 9±0.6 | 12±1.8 | 18±2 | 524±49 | 74±6 | 22±3 | 46% | 0.2±0.03 |
| **P21-24** (n=19) | 0.2±0.01 | 2.2±0.18 | | 9.1±0.6 | 16±1.5 | | 593±58 | 82±7 | | 54% | 0.5±0.09 |
| MNTB neuron without axon and dendrite (n=4) | 0.2±0.03 | | | 14±1.7 | | | 650±52 | | | | 0.91±0.1 |
| MNTB neuron with axon and without dendrite (n=5) | 0.15±0.02 | 1.9±0.18 | | 11.5±0.2 | 18±0.04 | | 767±122 | 97±14 | | | 0.55±0.12 |

**Figure 4.**
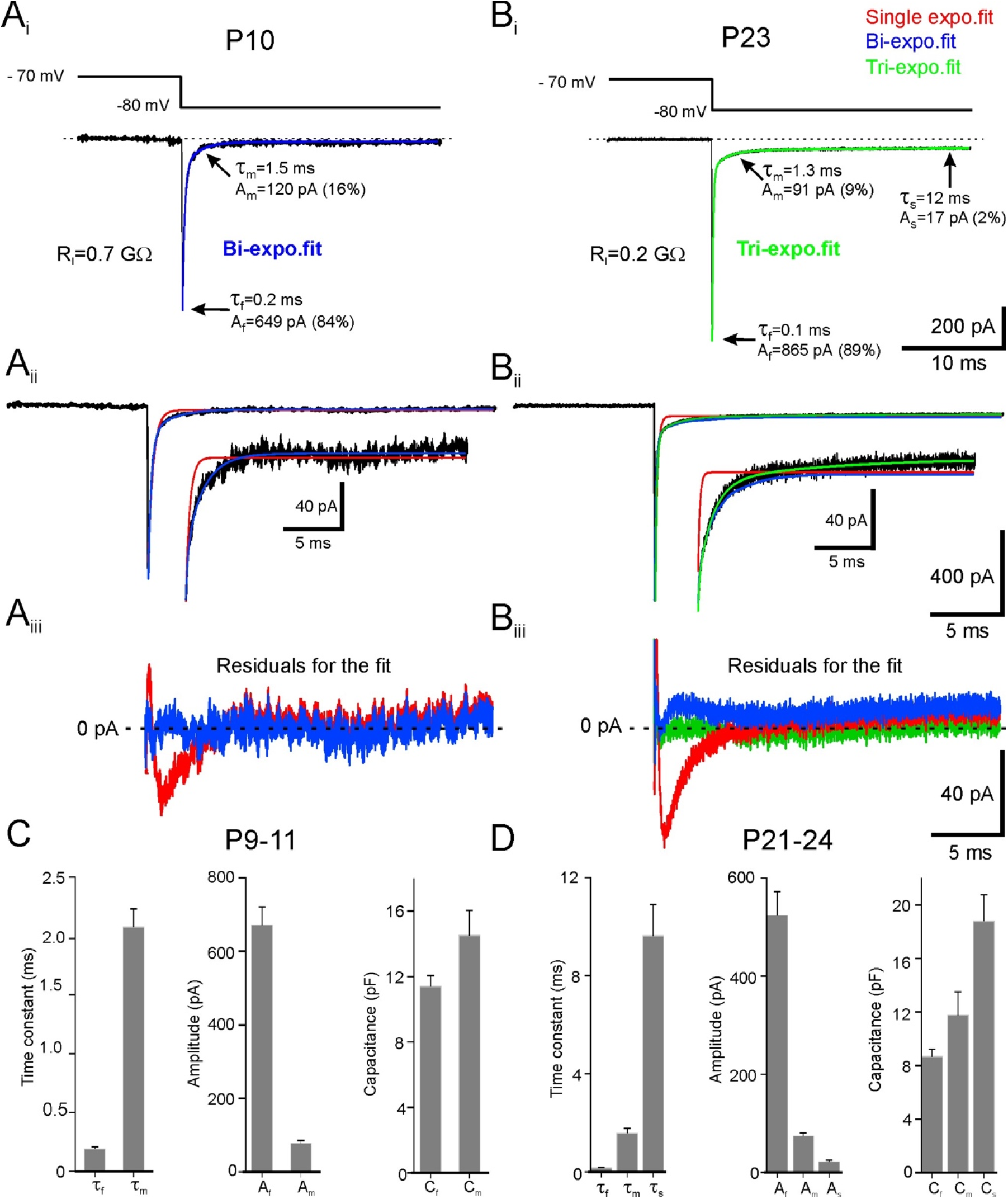
Voltage clamp responses of MNTB principal cells. Whole-cell recordings showing the passive properties of the MNTB neurons in response to 10 mV hyperpolarizing voltage step. (**A_i_**) Postnatal day 10 (P10) neuron showing a bi-exponential capacitive current decay (fit = blue curve) and a relatively small steady-state current during the voltage step. The membrane capacitances of the fast and medium components were 13 pF and 18 pF, respectively. (**B_i_**) Recording from a P23 neuron showing tri-exponential capacitive current decay (fit = green curve) and a relatively large steady-state current during the voltage step. The membrane capacitances of the fast, medium, and slow components were 9 pF, 12 pF, and 21 pF, respectively. (**A_ii_** and **B_ii_**) Capacitive currents from the same P10 and P23 neurons along with single exponential (red curve), bi-exponential (blue curve), and tri-exponential (green curve) fits. Inset shows the enlarged traces for clarity. (**A_iii_** and **B_iii_**) Difference between the measured capacitive currents and the fit with single, bi, and tri-exponential functions for the same P10 and P23 neurons. For the P10 neuron, a biexponential fit was sufficient to minimize residual current (blue trace), whereas for the P23 neuron a triexponential fit (green trace) was required. (**C**) Graph summarizing the mean decay time constants of the fast and medium components with their associated amplitudes and capacitance values (P9-11; n=26). (**D**) Graph summarizing the mean decay time constants of the fast, medium, and slow components with their associated amplitudes, and capacitance values (P21-24; n=16). Values are expressed as mean ± SEM.

From **Table 2** we see that the estimated C_m_ values for the P21-24 group of cells with three compartments are C_f_ = 9 pF and C_med_ = 12 pF and C_s_ = 18 pF, so that the total C_m_ = 39 pF. This is close to the estimates obtained with current clamp (45 pF; **Fig. 3E**) and SBEM (Table 1). However, do the three separate values of C_f_, C_med_ and C_s_ reflect the three distinct compartments of cell soma, dendrites and/or axon?

### Developmental changes in dendritic length

To determine the relationship between developmental changes in morphology and passive membrane properties we performed confocal reconstructions of MNTB neurons filled through the patch pipette with Alexa 555 dye (**Figure 5**). The size of the soma was measured as the geometric mean of major and minor axes and did not change significantly (16 ± 1.2 μm at P9-11 *vs*. 17 ± 1.6 μm at P21-24). Likewise, the length of the axon (measured by tracing in the confocal field of views) within the brainstem slices was not significantly different for the two age groups (172 ± 13 μm vs 203 ± 25 μm). At P9-11, MNTB neurons exhibited one or two very small dendritic stubs with an average length of 31±10 μm as shown in **Figure 5A** (see also Sätzler et al., 2002). By P21-24, the dendrites had grown significantly to 80 ± 17 μm (p=0.03; **Figure 5B**) and often bifurcated. We conclude that the major morphological difference likely to explain the changes in passive neuronal properties is the elaboration of dendritic structures that increase 2.6-fold in length between P9-11 and P21-24 as summarized in **Figure 5C**.

**Figure 5.**
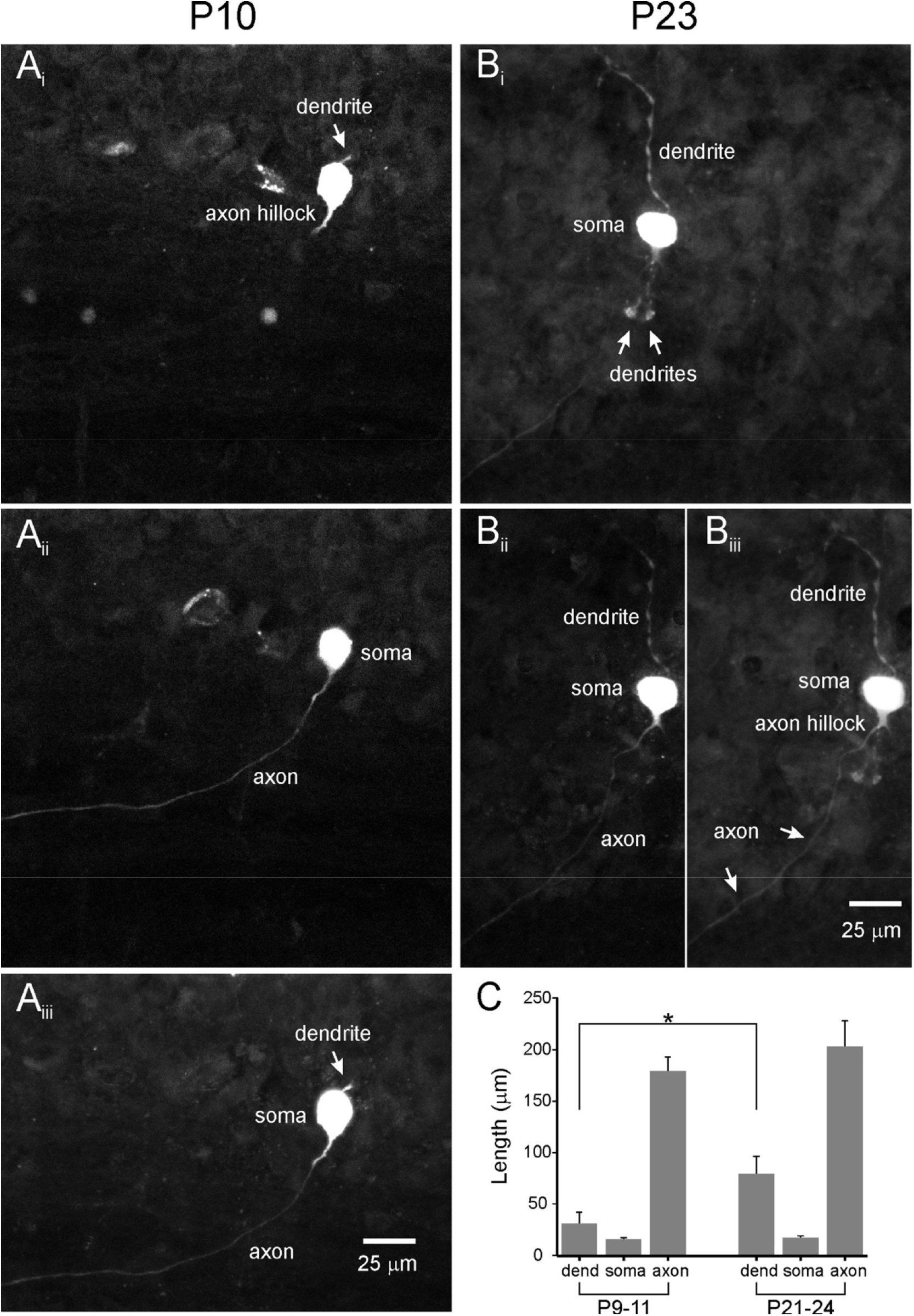
Confocal fluorescence images of MNTB neurons filled with Alexa fluor 555. (**A_i-ii_**) Fluorescence images showing the morphology of a P10 MNTB neuron (2 different frames). (**B_i-ii_**) Fluorescence images showing the morphology of a P23 MNTB neuron (2 different frames). (**A_iii_** and **B_iii_**) Maximum intensity projection of 10 image planes showing the morphology of the same P10 and P23 MNTB neurons. The P23 neuron shows comparatively longer and bifurcated dendrites than P10. (**C**) Graph summarizing the mean dendritic length, diameter of the soma, and axonal length (traced in the field of view) of P9-11 (n=11) and P21-24 (n=7) MNTB neurons. Asterisk indicate a significant difference in length (p=0.03).

### Membrane compartments and passive cell properties

We occasionally observed neurons with mono-exponential decay of their capacitive transients (4 of 44 cells at P9-17). Imaging showed that these neurons lacked both dendrites and axon, presumably lost during the brain slicing procedure. **Figure 6A** shows a cell exhibiting a mono-exponential decay. In such cells, the time constant was 0.2 ± 0.03 ms (n = 4), a value very similar to the fast component of decay in intact cells (p=0.23; Fig. 4B). The integral of the capacitive current divided by the voltage step gives the associated capacitance which was 14.0 ± 1.7 pF. Of all neurons tested, the mono-exponential decay group had the highest input resistance (0.9 ± 0.1 GΩ; **Figure 6D** & Table 2), indicating that the axon and dendrites make substantial contributions to input conductance. Somata of the MNTB neuron were ellipsoid. Based on the measurements of major and minor axis, the surface area of MNTB neurons is about 970 μm^2^ and the predicted membrane capacitance (assuming a smooth surface as demonstrated by EM; Fig. 1) is 9.7 ± 1.7 pF (n = 4). This value is similar to the value which we obtained by fitting the capacitive current decay by using the formula: τ_f_ × A_f_/10 mV (9.0 ± 0.6 pF, n = 4; **Fig. 6D**). The somewhat larger value obtained by integrating the capacitance transient presumably reflects contributions from axon or dendrite stubs. We conclude that the fastest component of capacitive current decay largely reflects charging of the soma and proximal portions of the axon or dendrite. As mentioned above, neurons lacking processes (n=4) had a large R_I_ of about 0.9±0.1 GΩ. This value was higher than neurons with an axon (0.54±0.05 GΩ; n=55) or additional dendrites (0.2±0.02 GΩ; n=20). Thus, adding compartments increases the cellular surface area, and presumably leak channels, which in turn decreased the R_I_. The calculated access (series) resistance (from τ = C_m_ x R_a_) was 14.3 MΩ, in good agreement with the estimate from the patch clamp amplifier.

**Figure 6.**
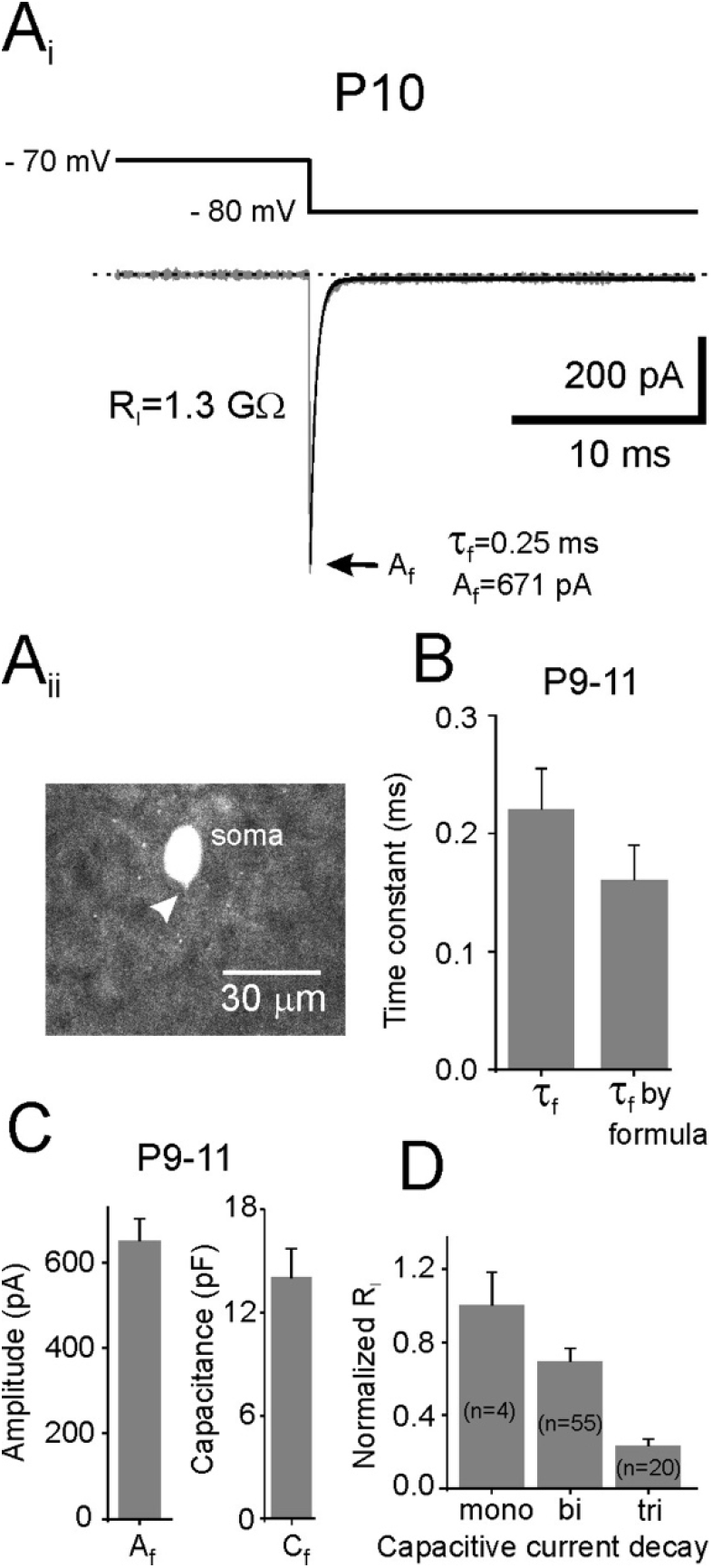
The passive properties of MNTB neurons which lack axon and dendrite. (**A_i_**) P10 neuron displaying capacitive current with mono-exponential decay and a small steady-state current upon −10 mV voltage step. (**A_ii_**) Fluorescence image of the representative neuron whose recording is shown above. The arrow head points to very small neurite process (either a cut axon or a small dendrite stub). (**B**) Bar graph summarizing the mean decay time constants of the capacitive current from neurons lacking any process. Both values (fitted and calculated using the membrane time constant formula) are similar (p=0.23). (**C**) Graph summarizing the mean amplitude and capacitance (P9-17; n=4). (**D**) Summary graph of all recorded cells showing the correlation of capacitive current decay with input resistance (R_I_).

Given these results with single compartment cells, which lack axons and dendrites, we suggest that C_f_ values reflect mostly the soma. The P9-11 cells with two components suggest that that C_med_ reflects mostly the lightly myelinated axon. Thus, for cells with three compartments, C_s_ may reflect mostly the dendrites. However, there is some uncertainty on how much one can separate axon and dendritic capacitance with these methods given that they have similar calibers initially as they leave the soma (1.5 μm for dendrites and ~ 1.0 μm for the myelinated axon). However, at P21-24 myelin will greatly reduce the axonal capacitance so it may have very little contribution to the overall C_m_ of the MNTB cell. We thus propose that for the P21-24 group of MNTB cells with three compartments the C_f_ = 9 pF reflects mostly the soma, but C_med_ = 12 pF and C_s_ = 18 pF (Figure 4D and Table 2), probably reflect the 2 to 3 long and thin dendrites and to some unknown degree the axon.

### Influence of dendrites on signal transfer: EPSCs and EPSPs

Developmental changes in EPSC time course also affect the time course of the EPSP (Joshi and Wang, 2002; Taschenberger and von Gersdorff, 2000). In addition, an increase in potassium and I_h_ currents during development can also change the EPSP time course (Golding, 2012; Leão et al., 2010). To determine the relative importance of these factors, we measured EPSCs and EPSPs from the principal cells of the MNTB during afferent fiber stimulation. Typical examples (average of 5 sweeps at 0.1 Hz) are shown in **Figure 7A and 7B**. Postsynaptic action potentials were blocked by QX-314 in the patch pipette solution. EPSCs from P9-11 synapses exhibited a bi-exponential decay with time constants of 1.0 ± 0.1 ms (92% of current amplitude) and 7.8 ± 0.6 ms (n=14; **Fig. 7C**). At P21-24, EPSC decay was mono-exponential with a fast time constant of 0.5 ± 0.07 ms (n=8). The very fast kinetics of the EPSCs means that the voltage response is predicted to be determined largely by the passive membrane time constant (τ_M_ ~ 6-7 ms at P20 for 24°C; Fig. 3). Indeed, EPSPs from P9-11 neurons exhibited a bi-exponential decay with time constants of 5.5 ± 1.2 ms (64 % of decay) and 31.0 ± 4.7 ms (n=8; **Fig. 7D**), and a weighted time constant τ_wd_ = 14.7 ms (see Methods). By contrast, the EPSP decay at P21-24 was a bi-exponential function with much faster decay kinetics of 1.8 ± 0.5 ms (96% of decay) and 8.0 ± 2.23 ms (n=6), and thus τ_wd_ = 2.1 ms.

**Figure 7.**
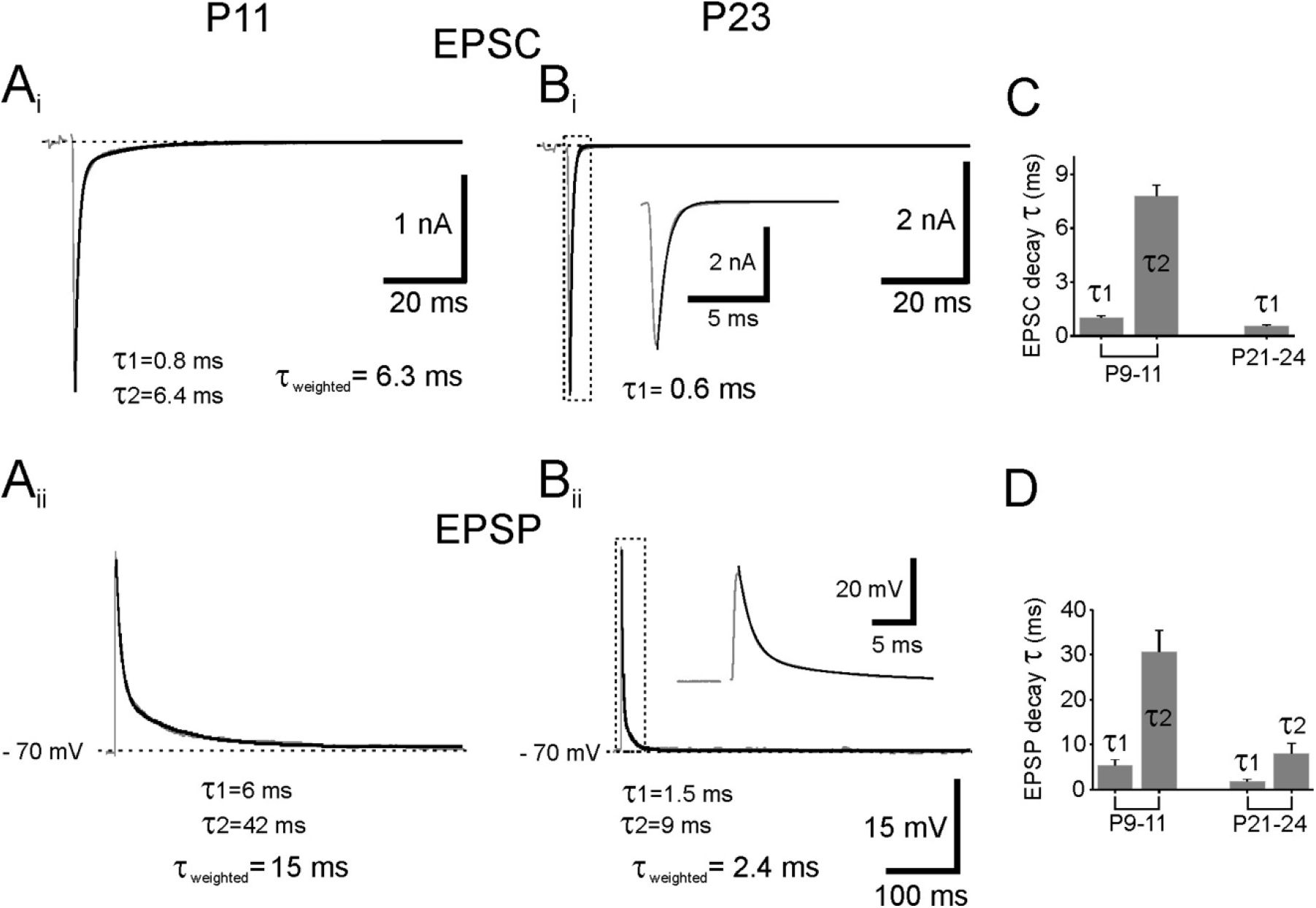
Decay kinetics of synaptic currents and synaptic potentials at younger and older ages during afferent fiber stimulation (0.1 Hz). (**A_i_** and **B_i_**) Evoked excitatory postsynaptic currents (EPSCs) of a P11 and a P23 MNTB neuron (grey trace) with their fitted decay shown in black. Inset shows enlarged trace for clarity. (**C**) Summary graph showing mean decay time constants of EPSCs at P9-11 (n=14) and at P21-24 (n=8). (**A_ii_** and **B_ii_**) Evoked excitatory postsynaptic potentials (EPSPs) of the same P11 and P23 MNTB neuron (grey trace) with their fitted decay kinetics (black). Inset shows enlarged trace for clarity. **(D**) Summary graph showing the mean decay time constants of EPSPs at P9-11 (n=8) and at P21-24 (n=6).

To examine the importance of voltage activated currents in a more functional context, we examined the decay time constants of large EPSCs and EPSPs generated at low frequency (**Fig. 7**) and the depressed EPSCs and EPSPs recorded at the end of a 100 Hz train of afferent fiber stimulation (**Figure 8**; see also Taschenberger and von Gersdorff, 2000). The slow decay of the last EPSC in **Figure 8Ai** may be due to glutamate pooling in the synaptic cleft of the P11 calyx synapse and a lower expression of glutamate transporters (Renden et al., 2005), whereas the faster decay of the last EPSC at P23 (**Figure 8Bi**) may be due to the fenestration of the calyx terminal into long finger-like stalks and swellings (Rowland et al., 2000).

**Figure 8.**
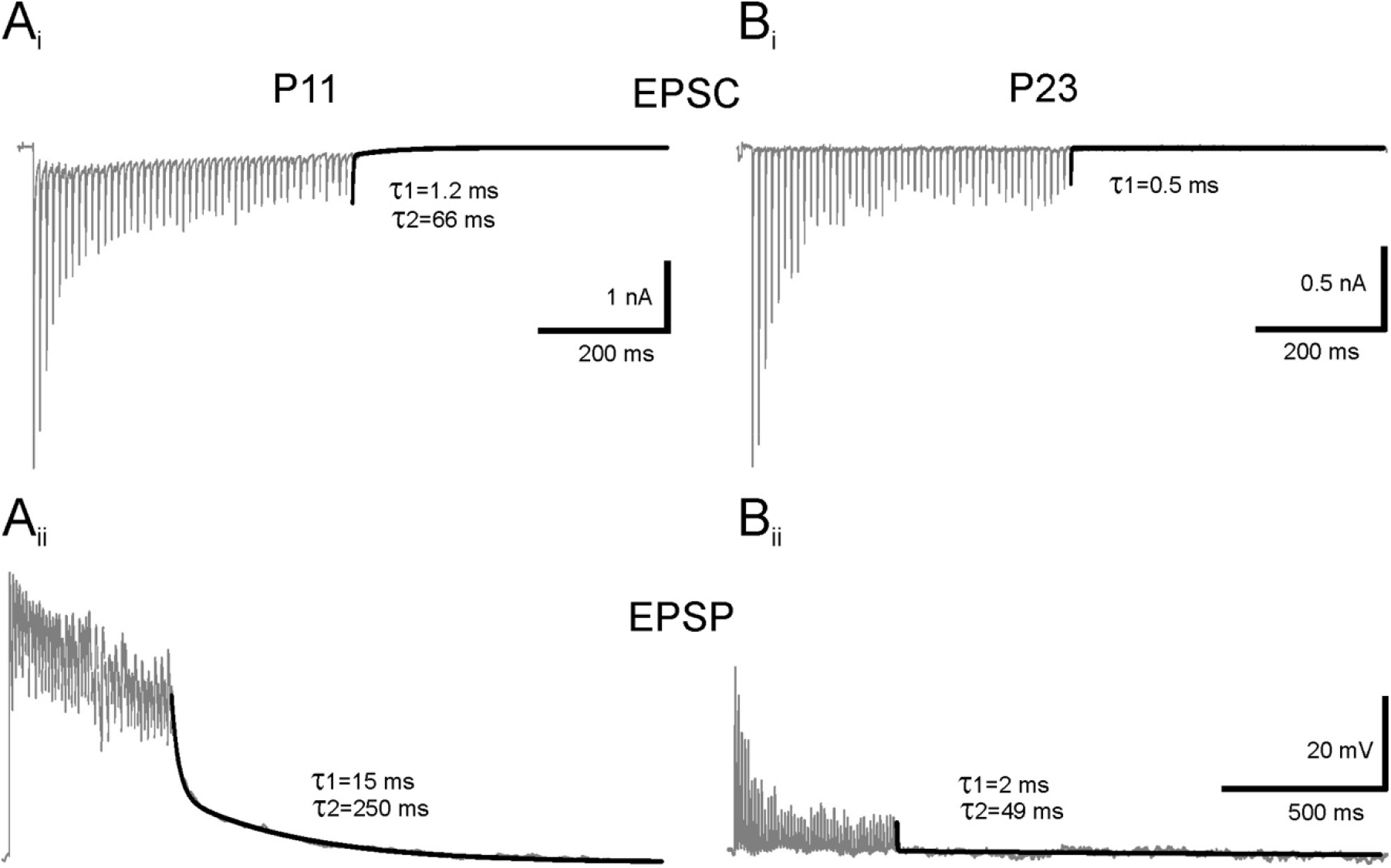
Decay kinetics of synaptic currents and synaptic potentials in MNTB neurons during 100 Hz train stimulation (50 stimuli). (**A_i_** and **B_i_**) Evoked EPSCs (grey traces) of P11 and P23 MNTB neurons with their fitted decay kinetics (black). (**A_ii_** and **B_ii_**) Evoked EPSPs of the same P11 and P23 MNTB neuron (grey trace) shown above and their fitted decay kinetics (black). P23 neuron showing faster bi-exponential decay than P11 neuron.

**Figure 8Aii** shows EPSPs obtained with QX-314 in the patch pipette to block Na^+^ currents and AP spikes. At P11 a large depolarizing plateau is produced by the EPSP train. This is due to the summation of the slowly decaying EPSPs and activation of NMDA receptors (Taschenberger and von Gersdorff, 2000). At P23, after blocking postsynaptic APs, the fast decaying EPSPs do not summate (**Figure 8Bii**). These data indicated a slower decay of the small depressed EPSPs recorded at the end of the train compared to the single EPSPs shown in **Figure 7**. Similar results were observed in 4 recordings at P9-11 and 4 at P21-24. The dramatic differences shown here illustrate the importance of rapid EPSC decay and short membrane time constant for a rapid EPSP decay.

We also next determined whether the weighted time constant of EPSCs and EPSPs were correlated with the membrane resting input resistance (R_I_). **Figure 9A** shows the relationship between the weighted decay time constants of EPSCs with respect to R_I_. The weighted decay time constant of the EPSCs in younger (P9-11; n=12) and older (P21-24; n=9) MNTB neurons did not correlate with R_I_ (r=0.10 for P9-11 and r=0.001 for P21-24 neurons). By contrast, the weighted decay time constant of the EPSPs showed a positive correlation with R_I_ (**Fig. 9B**). Moreover, P21-24 neurons displayed a steeper slope and exhibited a strong linear correlation (r=0.46 for P9-11 and r=0.82 for P21-24 neurons). We conclude that the observed changes in membrane passive properties during development have important consequences for the time course of EPSPs in MNTB neurons.

**Figure 9.**
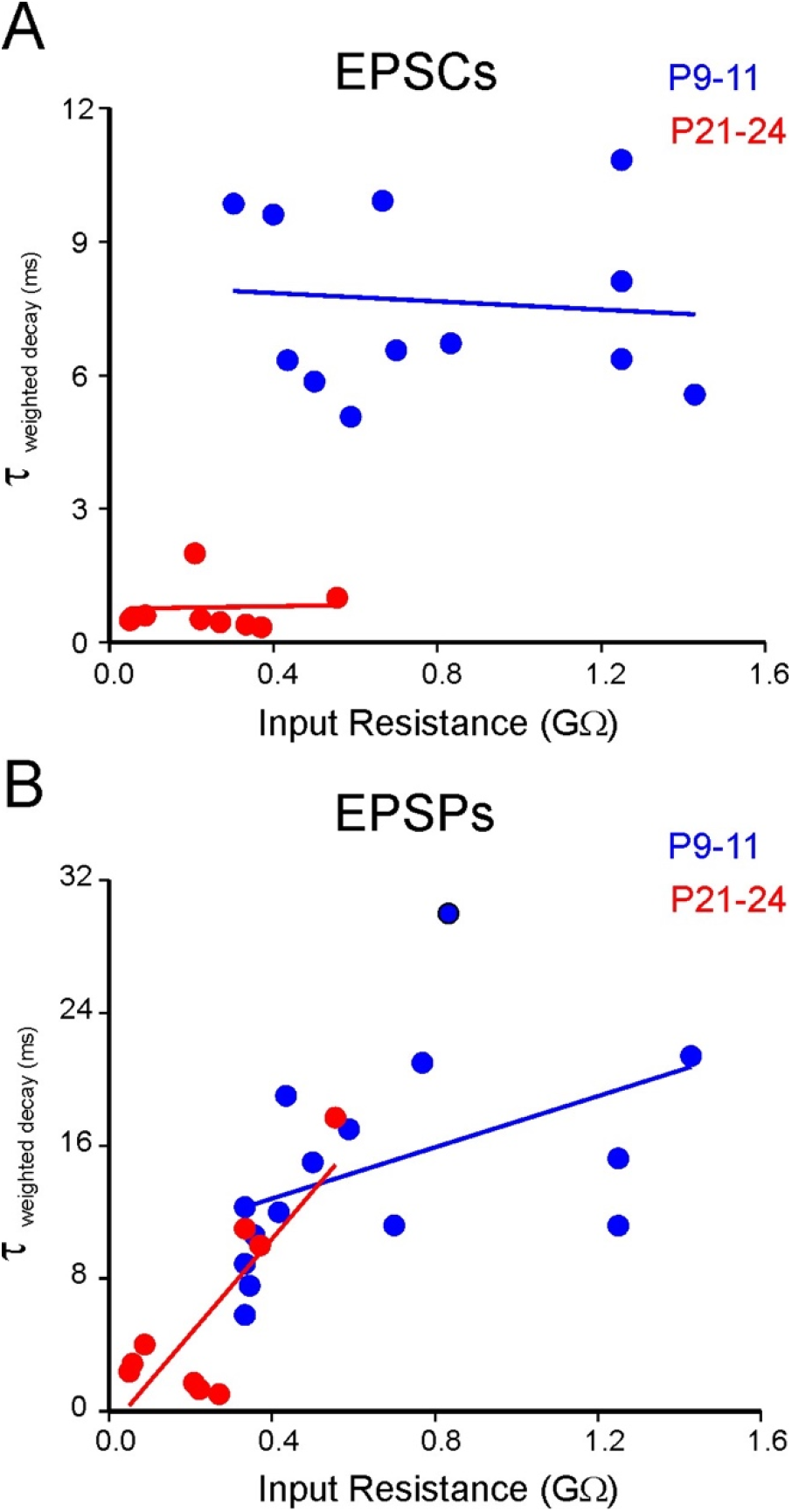
Graph summarizing the relationship of decay kinetics of synaptic currents and synaptic potentials with respect to R_I_. (**A**) Summary graph showing lack of correlation of weighted decay time constants of EPSCs and R_I_ of P9-11 (blue; n=12) and P21-24 (red; n=9) MNTB neurons. (**B)** Summary graph showing the correlation of weighted decay time constants of EPSPs and R_I_ of P9-11 (black; n=15) and P21-24 (red; n=9) MNTB neurons.

### Dendrites accelerate the EPSP decay

To further examine the correlation between the capacitive currents, R_I_, dendrites, and their synaptic currents and potentials, we examined the fraction of older neurons (P21-24; n=5) which displayed bi-exponential capacitive current. The morphology of these cells showed a prominent soma and a long axon, but little or no dendrites (perhaps lost during slicing; **Figure 10A)**. For these cells, the decay time constants of the capacitive currents were τ_f_ (0.15±0.02 ms) and τ_m_ (1.9±0.18 ms). These cells had a relatively large R_I_ = 0.55 ± 0.12 GΩ. We also checked the decay time constants of both the EPSCs and the EPSPs of these neurons. The weighted decay time constant of the EPSCs and EPSPs were 0.8±0.3 ms and 9.84±1.4 ms, respectively. We next compared these neurons which lacked dendrites (P21-24; n=5) to neurons with dendrites (P21-24; n=5). The weighted decay time constant of EPSC in neurons with dendrites was 0.5±0.03 ms which was not different from neurons without dendrites (p=0.21; Fig. 8B). However, the weighted EPSP decay time constant of neurons with dendrites was significantly faster (τ_wd_ = 1.85 ± 0.33 ms; p=0.0005; **Fig. 10B**). Thus, P21-24 neurons with dendrites and triphasic capacitive current decay have a 5-fold faster EPSP decay compared to neurons lacking dendrites and showing bi-phasic capacitive current decay (**Fig. 10C**). In fact, P21-24 neurons without dendrites had passive membrane properties and EPSP kinetics more similar to P9-11 neurons. We therefore suggest that it is the presence of dendrites that causes the major observed developmental changes in MNTB membrane properties and the fast decay of the EPSP.

**Figure 10.**
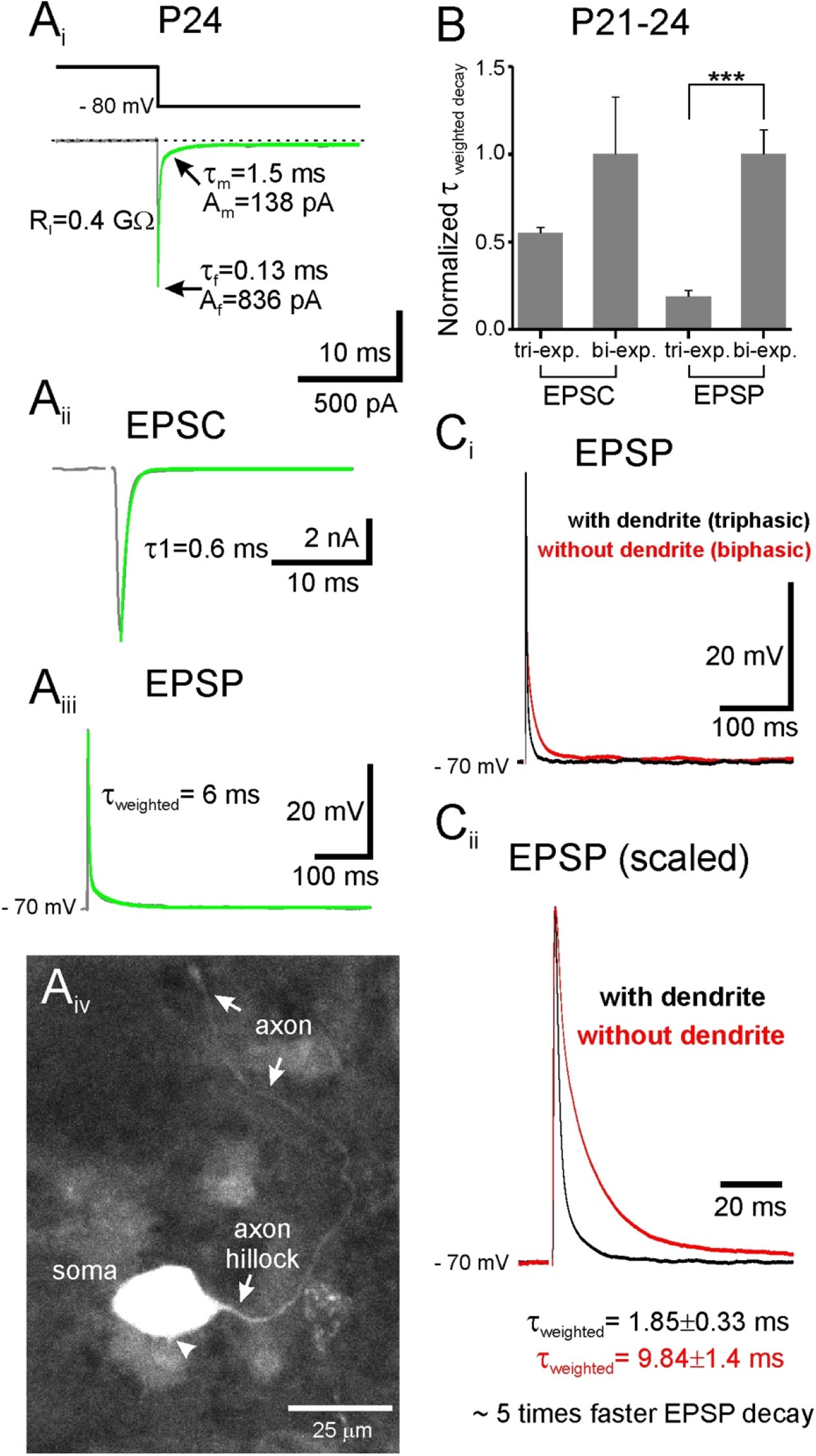
Passive properties of older MNTB neurons with axons and with or without long dendrites. (**A_i_**) P24 neuron gives a bi-exponential capacitive current decay (fit in green) and a relatively small steadystate current during the voltage step, indicating a high input resistance. (**A_ii_** and **A_iii_**) Evoked EPSCs and EPSPs of the same neuron with its associated decay kinetics. (**A_iv_**) Fluorescence image showing the morphology of a typical neuron which possesses an axon but lacks dendrites. The arrow points to the axon and the arrowhead to a putative dendrite stub. (**B**) Summary graph showing the relationship of triexponential and bi-exponential capacitive current decay with respect to EPSCs and EPSPs (P21-24; n=5 cells in each case). EPSPs significantly decay faster with tri-exponential capacitive current (p=0.0005). (**C_i_** and **C_ii_**) Averaged evoked EPSP from older neurons with dendrite (black; n=5) and without dendrite (red; n=5). Note that the neurons with dendrite EPSP decays ~ 5 times faster than neurons lacking dendrites.

### Computer modeling of EPSPs and leaky dendrites

In order to synthesize and interpret our data from voltage and current clamp measurements we generated a compartmental model of an MNTB neuron. For a point neuron (or a single and uniform spherical compartment), the membrane time constant is given by τ_m_ = C_m_ /G_m_, the ratio of membrane capacitance (in F/cm^2^) to resting leak conductance (in S/cm^2^). Thus, increasing the total membrane area will not change the membrane time constant. As an example, a point-neuron model with a standard C_m_ = 10 fF/μm^2^ = 1 μF/cm^2^ and G_m_= 200 μS/cm^2^ has a single exponential time constant τ_m_ = 5 ms and a synaptic current I = 5 nA generates a peak EPSP amplitude of 10 mV, if the resting membrane conductance is g_m_ = 0.5 μS (V_m_ = I/g_m_; see Golding, 2012).

How does EPSP decay change after an increase in dendrite length and dendritic conductance in a non-point neuron? To gain further insights on the role of dendritic length and leak conductance on EPSP decay, we generated passive compartmental MNTB cell models that consisted of a single dendrite of various lengths, a soma and a thin axon (**Figure 11**). We applied single EPSPs to the soma based on the model of Graham et al (2001) and Leão et al. (2008). Modeling parameters and decay values are listed in **Table 3**. EPSPs decays were fit with double exponentials. When the cell was implemented with a 10 μm long dendrite and a leak conductance (*g_L_*) of 62.5 μS/cm^2^, the weighted decay time constant was 11.6 ms (**Figure 11Ai**). After increasing *g_L_* to 500 μS/cm^2^, maintaining the dendrite length equal to 10 μm, the EPSP weighted decay time-constant was 9.7 ms, Using an 80 μm-dendrite, the weighted decay time was 11 ms for *g_L_*=62.5 μS/cm^2^ and 6.5 ms for *g_L_*=500 μS/cm^2^ (**Figure 11Aiv**). **Figure 11B** shows that without dendrites the EPSP decays more like a single exponential and does not match the clear double exponential decay we observed in Figure 3C with short current injections. **Figure 11C** shows the expected result that just adding longer dendrites with a constant total conductance *g_L_* = 250 μS/cm^2^ does not change EPSP decay kinetics. Thus, cells with longer dendrites that increase the leak conductance produce faster double-exponential decay kinetics for simulated EPSPs.

**Figure 11.**
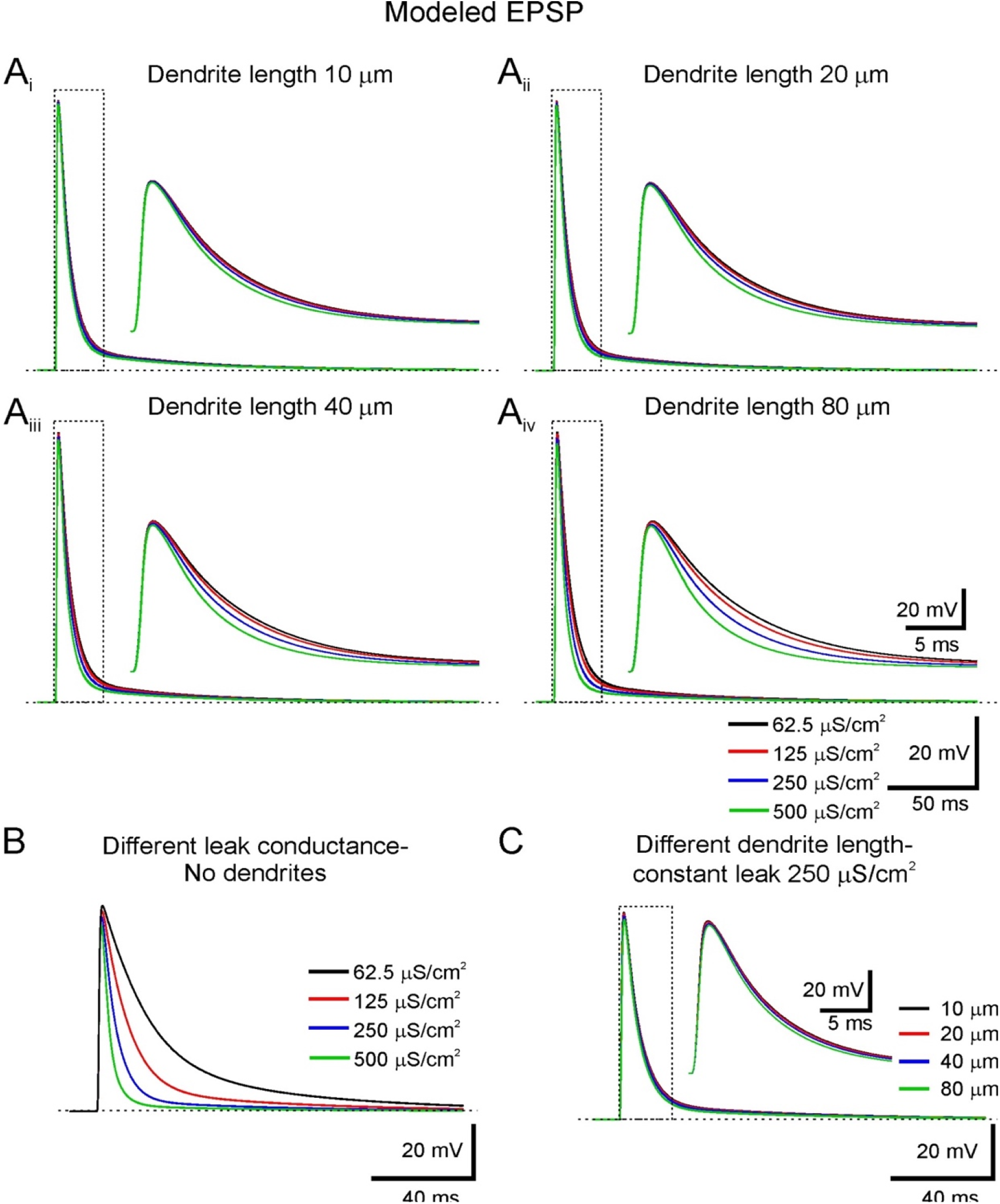
Model neuron response to simulated single calyceal EPSPs. (**A**) Simulated EPSPs response of a modeled MNTB neuron with different dendrite lengths (**A_i-ii-iii-iv_**: 10, 20, 40, 80 μm) and with different leak conductance on the dendrite (black: 62.5 μS/cm^2^, red: 125 μS/cm^2^, blue: 250 μS/cm^2^, green: 500 μS/cm^2^). (**B**) Simulated EPSPs with different leak conductance, but without dendrite (black: 62.5 μS/cm^2^, red: 125 μS/cm^2^, blue: 250 μS/cm^2^, green: 500 μS/cm^2^). (**C**) Simulated EPSPs with different dendrite lengths and with constant leak conductance (250 μS/cm^2^). Note that the simulated EPSPs decay faster with larger leak and longer dendrites, which simulates the behavior of the experimental results.

**Table 3.** Computer simulated EPSP weighted decay time constants using different dendrite lengths and leak conductance.

| Leak conductance | Dendrite length-Weighted decay $\tau$ | | | | |
| --- | --- | --- | --- | --- | --- |
| | 10 $\mu\text{m}$ | 20 $\mu\text{m}$ | 40 $\mu\text{m}$ | 80 $\mu\text{m}$ | No dendrite |
| 62.5 $\mu\text{S}/\text{cm}^2$ | 11.61 ms | 11.24 ms | 11.21 ms | 11.2 ms | 22.2 ms |
| 125 $\mu\text{S}/\text{cm}^2$ | 11.28 ms | 11.04 ms | 10.75 ms | 10.4 ms | 14.35 ms |
| 250 $\mu\text{S}/\text{cm}^2$ | 10.74 ms | 10.26 ms | 9.41 ms | 8.63 ms | 10.9 ms |
| 500 $\mu\text{S}/\text{cm}^2$ | 9.7 ms | 9.0 ms | 8.0 ms | 6.5 ms | 4.9 ms |

Note that EPSP amplitudes are normalized in **Figure 11** to better compare the EPSP decay. Of course, a large leak conductance, mediated for example by low threshold K^+^ channels, will also decrease the amplitude of the EPSP because V_m_ = I/g_m_ (Golding, 2012). Thus, if a large leak conductances are necessary to produce a fast EPSP decay, a large EPSC is also necessary to maintain a super-threshold EPSP amplitude. We conclude that an increase in leak conductance will decrease membrane time constant and will speed up EPSPs provided the membrane time constant is slower than the synaptic conductance. This is indeed the case for the very fast and large AMPA-receptor-mediated EPSC of the calyx of Held synapse.

## Discussion

Our results show that the principal neurons of the MNTB undergo dramatic morphological changes during the first three postnatal weeks. The *raison de être* for the development of long and thin dendrites in MNTB neurons has remined mysterious given that the major excitatory input resides in the soma. In order to assess the contribution of dendritic properties in shaping the evoked EPSP kinetics, we studied the MNTB neurons with and without dendrites, which were presumably severed during the slicing procedure. We found that EPSP decay was faster in older neurons that had longer dendrites and a decreased input resistance. Dendrites may thus play an important role in determining the synaptic integration of auditory signals in the MNTB.

### MNTB neurons have long and thin dendrites

Dendrites are the main sites of synaptic input for most neurons and they play both active and passive roles in synaptic integration. While MNTB neurons receive both excitatory and inhibitory synaptic inputs onto the soma, the presence of functional dendritic inputs is not clear (Smith et al., 1998; Hamann et al., 2003; Leao et al., 2004). Electron microscopic studies describe some non-calyceal small bouton type terminals on the cell body and dendrites (Lenn and Reese, 1966; Elezgarai et al., 2003). Anatomical evidence also suggests that the calyx of Held can make occasional contact with proximal dendrites (Morest, 1973; Rowland et al., 2000). The dendrites of MNTB neurons are also enriched with Na^+^ and K^+^ channels, which probably play a major role in promoting and adapting the firing pattern of the MNTB cell (Perney and Kaczmarek, 1997; Elezgarai et al., 2003; Leão et al., 2008).

Our SBEM results reveal that P30 mouse MNTB cells have 1 to 3 thin dendrites (diameter ~ 1.5 microns). The dendrites bifurcated into 2-3 thin branches and spanned an overall distance of about 80 to 200 microns. They receive sparse synaptic bouton input (Figure 2). Our confocal imaging with Alexa dye revealed that P21-24 MNTB neurons also have similar thin and long dendrites that extended for 80 microns. However, P9-11 neurons had 1 or 2 shorter dendrites that extended for only about 25 microns. The overall morphology of the P21-24 cells was very similar to the morphology revealed by SBEM for the P30 cells. Indeed, our patch clamp recordings reveal an overall C_m_ of about 39 pF for P21-24 cells, whereas the total surface area of the P30 cells measured with SBEM is equivalent to about 33 to 41 pF, depending on various assumptions of cell shrinkage due to fixation and the membrane specific capacitance.

### Passive properties change during development

The majority of the younger neurons (P9-11) displayed biexponential capacitive current decay, whereas a major subset of more adult-like (>P21) neurons showed an additional third component. During the same time period, the input resistance declined with development (from about 0.43±0.04 GΩ at P11 to 0.20±0.02 GΩ at P23; p=0.003). The amount of current needed to elicit a single action potential increased during development (from 175 ± 9 pA for P11 to 200 ± 24 pA for P23; p=0.048). After measuring their passive properties, the neurons were imaged with confocal microscopy and the dendritic length was observed to double after mouse pup weaning (>P21) compared to young immature animals (~P9-11; pre-hearing), whereas the mean diameter of the soma and the axonal length remained constant (see Figure 4 and 5). This correlation of physiological properties and imaging suggests that the additional third capacitive current component observed during development may reflect the longer dendrites in older neurons.

### Why locate “leak” current and Na^+^ and K^+^ channels on long and thin dendrites?

Several different potassium channels are located on MNTB principal cells (Berntson and Walmsley, 2008; Brew and Forsythe 1995; Johnston et al., 2010). An action potential in the calyx of Held triggers glutamate release onto the soma of the MNTB principal cell. This generates a brief and large inward current that rapidly depolarizes the soma producing a fast rising EPSP that rapidly depolarizes the axon initial segment (AIS), which has a high density of Na^+^ channels (Leão et al., 2005). The depolarization then opens low threshold K_v_1.1/1.2 potassium channels in the AIS and high threshold K_v_3.1 potassium channels in the soma, dendrites and AIS (Johnston et al., 2010). Interestingly, K_v_3.1b is strongly concentrated in the dendrites of P16 and adult rat MNTB cells (Elezgahrai et al., 2003). This large I_K_ current quickly hyperpolarizes the soma back to resting membrane potentials via an outward flux of K^+^ ions. However, if these K_v_ channels were concentrated in the synaptic cleft or near the calyx of Held they may accumulate and depolarize the calyx membrane potential. This may be deleterious for a well-timed release of glutamate. Thus, localizing K_v_ channels to the dendrites and AIS may place them at a safe distance from the calyx, where they can produce a large I_K_ that does not influence the high frequency firing of the calyx nerve terminal. Indeed, K_v_3.1b potassium channels are excluded from the synaptic cleft of the calyx of Held (Elezgahai et al. 2003).

MNTB neurons have a low density of Na^+^ channels in their soma (Leão et al., 2005). We speculate that excluding Na^+^ channels from the soma and synaptic cleft avoids a potential reduction in the Na^+^ ion concentration in the narrow synaptic cleft, which has a narrow 20 nm width and where AMPA receptors are located. Importantly, a low density of somatic Na^+^ channels has also been observed at other auditory brainstem neurons, where it allows the EPSP to have a faster decay (Yang et al., 2016). Surprisingly, the dendrites of the MNTB cell have a high density of Na^+^ channels (Leão et al., 2008). These may boost the EPSP amplitude, but also slow its decay (Leão et al., 2008; Yang et al., 2016; Ceballos et al., 2017). Moreover, MNTB neurons also have Slick and Slack Na^+^-dependent K^+^ channels that regulate their high-frequency firing (Yang et al., 2007; Brown and Kaczmarek, 2011). If these K^+^ channels are located in the dendrite, next to Na^+^ channels, they also may play a critical role in speeding up the EPSP decay.

### Firing spikes at high rates in the AIS

The large depolarization of the somatic EPSC produces a rapid opening of Na^+^ channels in the long (10 to 15 micron) and thin unmyelinated axon initial segment (AIS; see Figure 1B), which has a high density of Na^+^ channels (Leão et al., 2005). Our SBEM reveals that the C_m_ of the AIS is only 1.1 pF. A large EPSP can thus quickly bring this small area to threshold with the opening of relatively few Na channels. A combination of local I_KLT_ and I_KHT_ (K_v_3.1) type channels then produces a rapid outward flux of K^+^ ions that quickly brings the membrane potential back to resting levels. The high density of Na channels present in the AIS allows for a large surplus of available Na channels for high frequency firing (Madeja, 2000). This large surplus allows the postsynaptic MNTB cell to fire spikes at rates as high as 800 Hz for extended periods of time (Taschenberger and von Gersdorff, 2000).

### Computer modeling: Leaky dendrites speed up the EPSP decay

The role of the dendrite in determining integrative properties of MNTB cells was further investigated by a simple computational model of a passive neuron. Our model consisted of a soma, axon and dendrites of various lengths (or devoid of dendrite) and various levels of leakage conductance. The simulations demonstrate that dendrites in MNTB neurons can minimize temporal summation of synaptic currents by shortening the decay of EPSPs. A longer dendrite augments considerably the membrane area and overall C_m_, which would slow EPSP decay if it had a small or no leak conductance. However, they also serve as extra remote sites for locating leak potassium channels or low threshold potassium channels. Hence, the dendrite can serve as a sink for the calyceal EPSPs generated at the soma, accelerating the decay of the EPSP. Our model also showed that the dendritic effect on EPSP decay is mostly attributed by the lowering of input resistance rather than changes in surface area (or capacitance) due to longer dendrites.

Our main experimental observation is that MNTB neurons that lack dendrites have a slower decay of the EPSP (Figure 10C). These results uncovered a dendrite-mediated speeding for the EPSPs for more mature MNTB neurons that is analogous to what has been proposed for interneuron axons in the cerebellum (axonal speeding) by Mejia-Gervacio et al. (2007). These authors observed a slower decay of the EPSP when they cut the interneuron axon with a laser. Our computer simulations also suggest that longer dendrites with a larger leak conductance produce faster EPSP decay kinetics (Figure 11). In summary, our results suggest that the thin and long dendrites of MNTB neurons provide a fast current sink that accelerates synaptic potentials. This reduces EPSP summation during long trains of repetitive high frequency afferent fiber stimulation.

## Acknowledgements

This work was supported by RO1 NIDCD grants to (HvG) and (GS) and CNPq Brazil (RNL).

